# Transgenic Targeting of *Fcrls* Creates a Highly Efficient Constitutively Active Microglia Cre Line with Differentiated Specificity

**DOI:** 10.1101/2023.09.30.559914

**Authors:** Tobias Kaiser, Jordan Dattero, Liang Li, Mandy Chen, Minqing Jiang, Andrew Harrahill, Oleg Butovsky, Guoping Feng

## Abstract

Microglia carry out important functions as the resident macrophages of the brain. To study their role in health and disease, the research community needs tools to genetically modify them with maximum completeness in a manner that distinguishes them from closely related cell-types, such as monocytes. While currently available tamoxifen-inducible CreERT2 lines are able to achieve the differentiation from other cells, the field needs improved and publicly available constitutively active Cre lines, especially ones with favorable efficiency and specificity profiles for studies where high recombination efficiency is imperative and where tamoxifen administration is contraindicated. Here, we leverage the microglia-specific *Fcrls* gene to generate mice expressing Cre. Using genomic methods, we show correct positioning of the transgene and intact microglia homeostasis in *Fcrls-2A-Cre* mice. Crossing *Fcrls-2A-Cre* mice to four different reporters, we demonstrate highly efficient recombination in microglia across differentially sensitive loxP alleles in different genomic contexts, indicating robust applicability of the line. Further, we show that microglia recombine a loxP reporter during early embryonic development, supporting the use of the line for developmental studies. Finally, using immunofluorescence and flow cytometry, we reveal that most border associated macrophages (BAMs) are also targeted whereas only few liver and spleen macrophages and virtually no white blood cell subsets exhibit Cre activity, distinguishing this line from another publicly available Cre line, *Cx3cr1-Cre*^*M*^ (MMRRC). *Fcrls-2A-Cre* mice are immediately available (JAX Stock #036591) and serve as a valuable addition to the community’s microglia toolbox by providing highly efficient constitutive Cre activity with excellent specificity, particularly for studies where tamoxifen administration is undesirable.

**Significance Statement:** The microglia toolbox is continuously growing with more transgenic lines and most recently even viral tools becoming available. When selecting a Cre driver line, investigators must weigh relative strengths and weaknesses of available lines and carefully make the best choice for their given application. These tradeoffs include (1) availability and ease of employment, (2) chromosomal positioning of Cre with respect to the floxed allele (should not be on the same chromosome for conditional knockout studies), (3) activity level of a given Cre line and thus completeness of recombination across the microglia population, (4) specificity with respect to acceptable off-target cell types and tissues, (5) temporal aspects including earliest onset of Cre expression or inducibility, (6) robustness in disease contexts, and (7) potential perturbation of microglia homeostasis through Cre itself or disruption of the targeting locus. When selecting a mouse line, it is evident that there may not be a one-size-fits all solution but an application-based preference and choice from the diverse repertoire of microglia tools. *Fcrls-2A-Cre* mice are an excellent addition to this toolbox.

## Introduction

Microglia are the resident macrophages of the brain (Lawson et al., 1990). Functionally, their phagocytic and secretory activities play a role in neurogenesis, development of neuronal connectivity, myelin maintenance, and survival of neurons (Stevens et al., 2007; Sierra et al., 2010; Paolicelli et al., 2011; Schafer et al., 2012; Ueno et al., 2013; Hagemeyer et al., 2017; Wlodarczyk et al., 2017; Weinhard et al., 2018). In addition, microglia react to perturbations such as vascular injury, multiple sclerosis lesions, and neurodegeneration (Itagaki et al., 1989; Davalos et al., 2005; Ransohoff, 2016; Aguzzi and Zhu, 2017; Keren-Shaul et al., 2017; Mathys et al., 2017; O’Loughlin et al., 2018). Dissecting the role of microglia in these processes hinges on mouse lines that enable conditional gene manipulation in microglia. Currently, several transgenic lines are available for constitutive (Cre) or inducible (CreERT2) deletion of floxed sequences. For Cre lines, loci harnessed include *Tie2, Csf1r, Lyz2, Cx3cr1, Cx3cr1/Sall1* (Clausen et al., 1999; Kisanuki et al., 2001 p.2; Ferron and Vacher, 2005; Kim et al., 2006; Samokhvalov et al., 2007; Yona et al., 2013; Orthgiess et al., 2016 p.2) and, more recently, *Crybb1* (Brioschi et al., 2023). For CreERT2, the currently most faithful lines target *Tmem119, HexB, P2ry12*, or *Cx3cr1* (Parkhurst et al., 2013; Yona et al., 2013; Kaiser and Feng, 2019; Masuda et al., 2020; McKinsey et al., 2020). When using these lines to study microglia, there are two major considerations. First, currently available constitutively active Cre lines often recombine floxed alleles in additional major non-myeloid cell types, as well as unwanted myeloid cells, such as monocytes (Orthgiess et al., 2016; Haimon et al., 2018; Masuda et al., 2020). Second, the high fidelity CreERT2 lines require tamoxifen administration, which is a relative contraindication for developmental studies, studies of sex-specific effects and laboratories with limited access or facilities. Thus, there is a critical need in the field for a constitutively active Cre line that distinguishes microglia from other closely related peripheral and central myeloid cells such as blood monocytes as well as peripheral macrophages (Wieghofer and Prinz, 2016; Haimon et al., 2018).

Recently, profiling of genes enriched in microglia compared to other myeloid cells revealed several candidate genes for the generation of new Cre lines (Butovsky et al., 2014). Fc receptor-like S, scavenger receptor (*Fcrls*) is expressed across all microglia subsets based on single cell RNA sequencing studies (Hammond et al., 2019; Li et al., 2019). Adding to this, bulk RNA sequencing along development shows that *Fcrls* expression commences early during embryonic development (Mass et al., 2016). In contrast, *Fcrls* is not expressed in monocytes and other peripheral immune cell subsets based on data made available through the ImmGen database (ImmGen.org). Together, these data make *Fcrls* a suitable target locus for the generation of a microglia-specific Cre line. Here, we report the generation and characterization of an *Fcrls-2A-Cre* knock-in mouse line, wherein microglia express Cre recombinase. By crossing this newly generated mouse line to four different floxed reporter alleles, we demonstrate functional Cre activity in all microglia and a large subset of border associated macrophages in the brain (BAMs). Further, we demonstrate that Cre is inactive in astroglia, oligodendroglia and neurons, and modestly but non-negligibly active in liver and spleen macrophages. Most strikingly, we found virtually no Cre expression in white blood cells, unlike the best currently available constitutively active Cre line, *Cx1cr1-Cre*^*M*^. Thus, this new mouse line will expand the microglia research toolbox by adding a highly efficient constitutively active Cre line with differentiated specificity. This line is immediately available to investigators from JAX (Stock #036591).

## Results

### Generation of a Constitutively Active Microglia Cre Line (*Fcrls-2A-Cre)* using a CRISPR/Cas9-mediated Knock-in Strategy

Cre-driver lines and conditional alleles are critical in the quest to understand the roles of microglia in health and disease. Several lines utilizing Cre or its inducible form CreERT2 were previously generated and are now widely used in research (excellently reviewed by Wieghofer and Prinz, 2016; Bennett and Bennett, 2020; Miron and Priller, 2020; Dumas et al., 2021). While these lines have collectively enabled the field to make significant headway in understanding microglia biology, additional publicly available transgenic mouse lines would provide valuable additions to our mouse line repertoire if they efficiently targeted microglia while obviating the need for tamoxifen-mediated Cre induction and while avoiding targeting of monocytes. Amongst a group of recently discovered microglia-enriched genes, we initially chose *Tmem119* to generate a constitutive *Tmem119-2A-Cre* mouse line in addition to previously published EGFP- or CreERT2-expressing lines (unpublished data, Bennett et al., 2016; Kaiser and Feng, 2019). Unlike the EGFP and CreERT2 line which showed specificity for microglia, the constitute Cre mouse line displayed widespread Cre activity in brain endothelial cells and was discontinued (unpublished data). Further mining of public datasets suggested that microglia-enriched *Fcrls* is a suitable locus for Cre knock-in due to its high expression across all microglia subsets, early onset of expression, and complete absence of expression in monocytes (Butovsky et al., 2014; Hammond et al., 2019; Li et al., 2019, ImmGen.org). To generate *Fcrls-2A-Cre* mice, we created a double-strand DNA targeting vector containing *2A-Cre*, 1.5 kb homology arms, and an additional PAM (protospacer adjacent motif) point-mutation and injected mouse zygotes with a mixture of this targeting vector, Cas9 protein, and three crRNAs (Figure 1A-B). This design inserts *2A-Cre-STOP* into the STOP codon of *Fcrls*, resulting in a single transcript that encodes both *Fcrls* and *Cre* by means of ribosomal skipping. Implantation of the zygotes into surrogates resulted in 14 live births (F0 founders) which we subsequently screened and genotyped with primer pairs having binding sites outside the homology arms and inside the transgene to generate amplicon uniquely present in mice with correct knock-in (Figure 1A). We found four out of the 14 founders positive for the insert in the *Fcrls* locus and crossed them to C57BL/6J to generate F1 animals, which we also found to harbor the insert in the *Fcrls* locus (Figure 1C). Using Sanger sequencing, we verified intactness of the genomic sequence and the 3’UTR single-base change introduced to create the PAM mutation that prevented re-cutting by CRISPR/Cas (Figure 1D). Together, these data demonstrate that we successfully generated *Fcrls-2A-Cre* mice.

**Figure 1.**
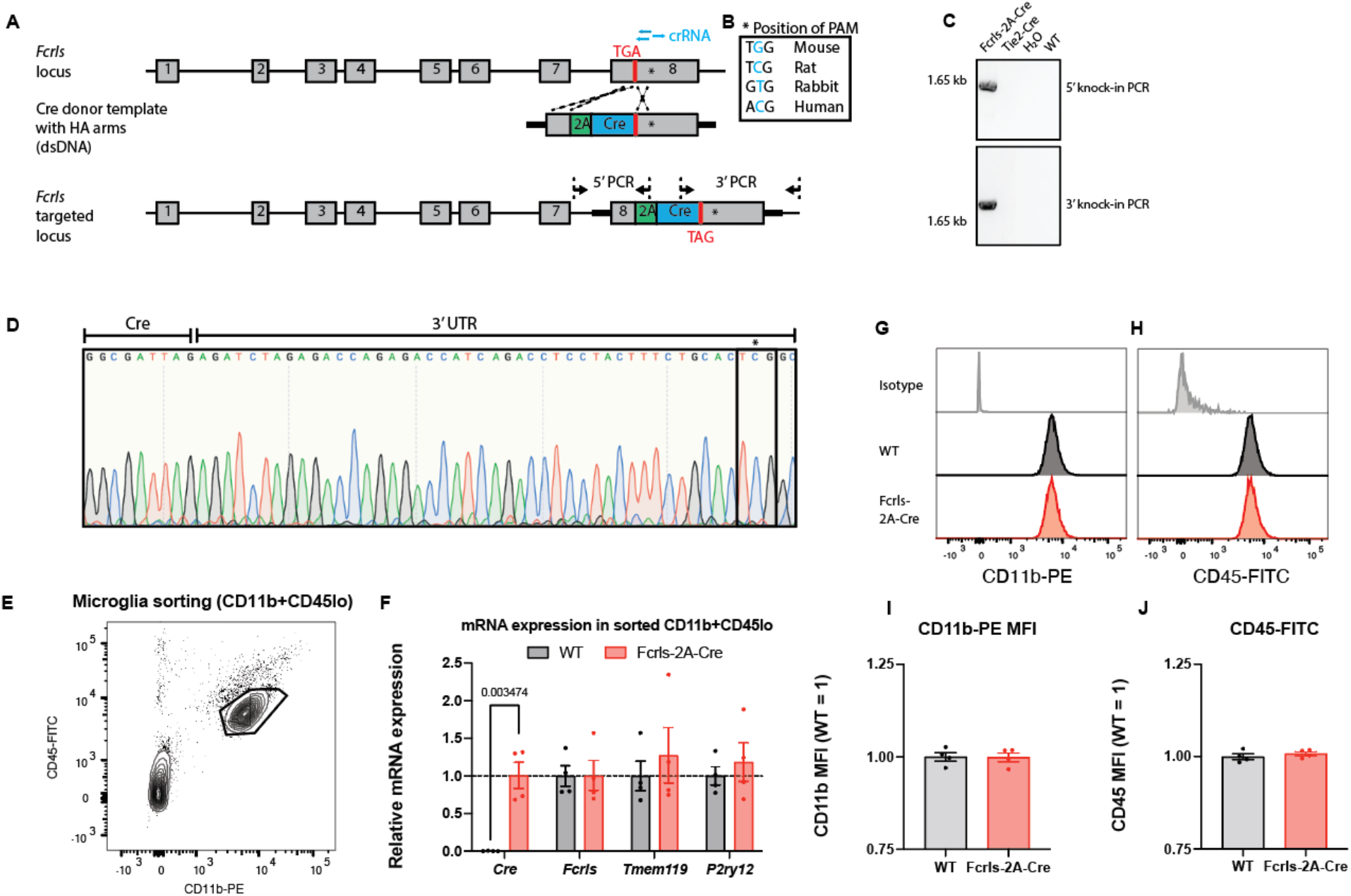
Generation of a Constitutively Active Microglia Cre Line Through Targeting of the *Fcrls* Locus Using CRISPR/Cas9. **A**, Schematic representation of murine *Fcrls* locus and knock-in approach to insert a *2A-Cre* cassette into the stop codon of *Fcrls* in exon 8 (not drawn to scale). Three crRNAs (blue bars) were selected to introduce double strand breaks at the stop codon and in the 3’ UTR. PCR primers to check for on-target insertion are indicated as arrows. **B**, The 3’ UTR-targeting sgRNA was selected such that nucleotide substitution required for silencing of the NGG PAM for CRISPR/Cas-mediated knock-in alters a non-conserved nucleotide (inset box). **C**, Representative agarose gel electrophoresis image of the indicated 5’ and 3’ PCR amplicons spanning the junctions of inserted *2A-Cre* transgene and target locus in founder animals. **D**, Sanger sequencing chromatogram of 3’ amplicon showing the G->C mutation in the founder animals. **E**, Representative flow cytometry density plot indicating gating of microglia for isolation (CD11b+CD45lo) in postnatal day 8 animals. **F**, RT-qPCR for *Cre* and *Fcrls* mRNA and two microglia homeostasis genes from sorted microglia. N=4 mice. Multiple t-tests with Benjamini, Krieger and Yekutieli correction for multiple testing. q=0.003 for *Cre*, others are not significant. **G-H**, Histogram plot for CD11b and CD45 expression on P8 microglia as gated in panel E. **I-J**, Quantification of CD11b and CD45 protein expression on P8 microglia. N=4 mice. Unpaired t-test. Not significant.

A drawback of using Cre to recombine floxed genes of interest and measure associated phenotypes is that the genomic Cre insertion or the Cre protein itself can affect cellular function. Specifically for microglia, Gosh and coworkers recently demonstrated that high Cre expression in *Cx3cr1-CreERT2* mice disrupts microglia homeostasis during early postnatal development (Sahasrabuddhe and Ghosh, 2022). To test if Fcrls-2A-Cre mice display similar abnormalities, we analyzed microglia from postnatal day 8 (P8) mice by flow cytometry and sorted them for subsequent RT-qPCR of homeostatic genes (Figure 1E). Unlike *Cx3cr1-CreERT2* P8 microglia analyzed by Sahasrabuddhe and Ghosh that displayed elevated CD11b and CD45 expression as well as disrupted homeostasis, we found microglia isolated from *Fcrls-2A-Cre* mice to be indistinguishable from WT microglia (Figure 1F-J). Together, these data suggest that microglia state in *Fcrls-2A-Cre* mice is unperturbed.

### *Fcrls-2A-Cre* Mice Effectively Recombine Floxed DNA in all Microglia and Most Border-Associated Macrophages (BAMs)

To investigate the activity of Cre recombinase, we crossed *Fcrls-2A-Cre* mice to Ai14 tdTomato mice, which report recombination with tdTomato fluorescence (Madisen et al., 2010). We prepared *Fcrls-2A-Cre*^*+/-*^; *Ai14*^*+/-*^ mice (short *Fcrls-2A-Cre; Ai14*, as only heterozygous mice were used throughout the study) for immunofluorescence staining of several regions. High power confocal micrographs of the cortex, hippocampus, and striatum showed that parenchymal IBA1-positive microglia expressed tdTomato (Figure 2A-C). All other CNS regions including the spinal cord showed robust tdTomato expression (data not shown). We quantified the confocal micrographs for completion (fraction of IBA1^+^ parenchymal microglia expressing tdTomato) and fidelity (fraction of tdTomato^+^ cells being parenchymal IBA1^+^ microglia) and found that parenchymal microglia were completely labeled and all labeled cells in the parenchyma were microglia (Figure 2D-E). In addition to parenchymal microglia, we examined border associated macrophages (BAMs), as these cells are closely related to microglia and at least a subset of them is known to express *Fcrls* in single-cell RNA-Sequencing analyses (Hammond et al., 2019; Li et al., 2019). Confocal micrographs of co-immunostained brain sections leveraging location of the cells showed that choroid plexus macrophages, meningeal macrophages, and perivascular macrophages all harbor recombined alleles to varying degrees, indicating Cre activity in a substantial fraction of these cells (Figure 2F-I). To further examine the specificity and to rule out any expression in non-myeloid cell types, we stained brain slices against S100B (astroglia), OLIG2 (oligodendrocyte lineage cells) and NEUN (neurons). Confocal micrographs revealed that none of the subsets expressed tdTomato (Figure 2 J-M).

**Figure 2.**
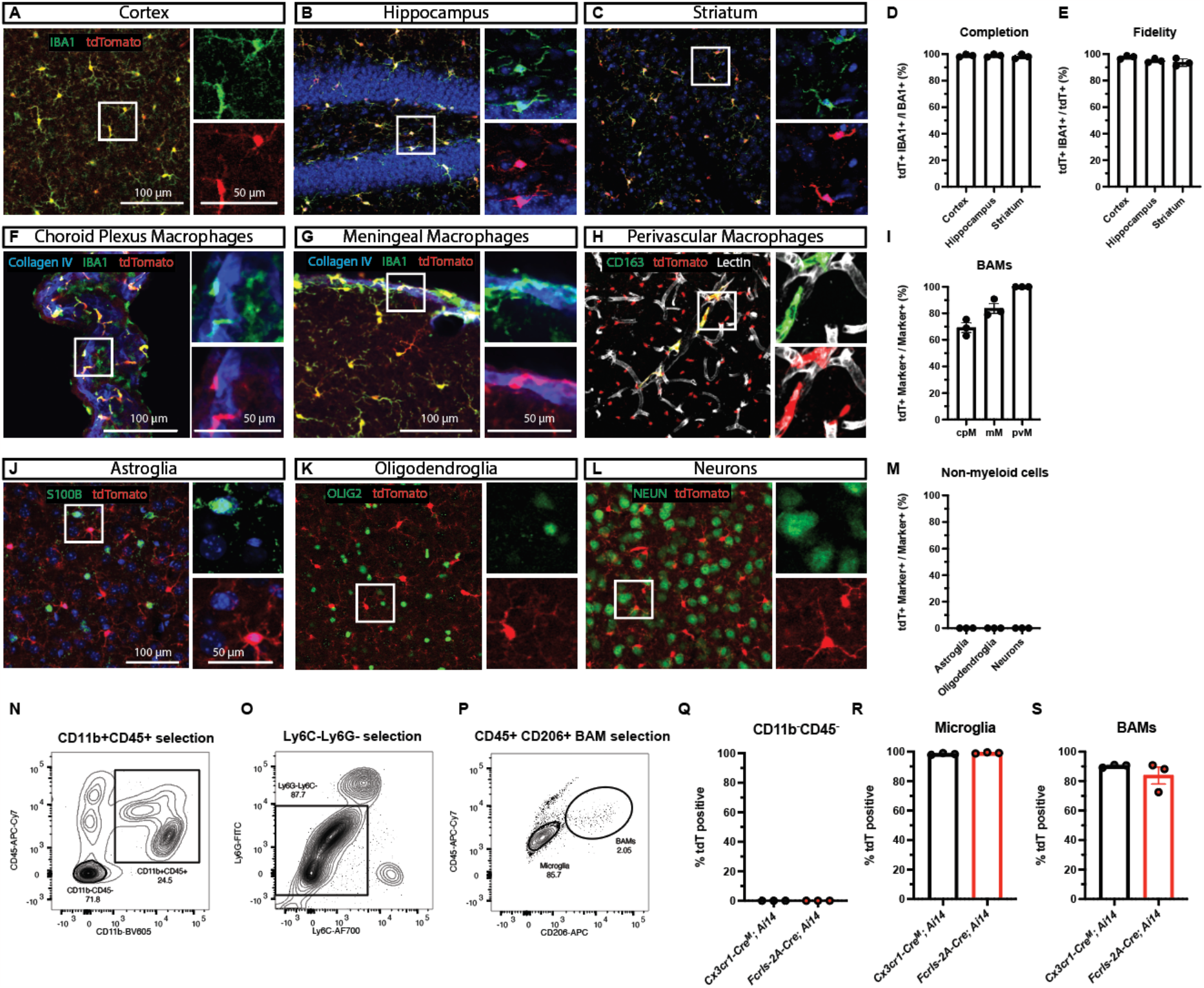
*Fcrls-A-Cre* Mice Effectively Recombine Floxed DNA in all Microglia and Most Border-Associated Macrophages (BAMs) **A-C**, Representative confocal micrographs of immunostained slices from *Fcrls-2A-Cre*^*+/-*^*;Ai14*^*+/-*^ mice showing tdTomato due to Cre recombination (red) and microglia (green) in cortex, hippocampus, and striatum. **D-E**, Quantification of the completion and fidelity of Cre activity in IBA1^+^ parenchymal macrophages in the brain. N=3 mice. **F-H**, Representative confocal micrographs of different border associated macrophage (BAM) subsets, including choroid plexus macrophages (green, anatomical, Collagen IV-adjacent), meningeal macrophages (green, Collagen IV-adjacent), and perivascular macrophages (green, CD163^+^, Lectin-adjacent). **I**, Quantification of *Fcrls-2A-Cre*-mediated recombination in BAM subsets. N=3 mice. **J-L**, Representative confocal micrographs of immunostaining for S100B (green, astroglia), OLIG2 (green, oligodendroglia), and NEUN (green, neurons) in slices from *Fcrls-2A-Cre*^*+/-*^*;Ai14*^*+/-*^ mice. **M**, Quantification of *Fcrls-2A-Cre*-mediated recombination in non-myeloid brain cells (astroglia, oligodendroglia, neurons). N=3 mice. **N-P**, Representative flow cytometry density plots showing gating of microglia (CD11b^+^CD45^lo^Ly6C^-^Ly6G^-^CD206-) and BAMs (CD11b^+^CD45^+^Ly6C^-^Ly6G^-^CD206^+^). **Q-S**, Quantification of *Fcrls-2A-Cre*- or *Cx3cr1-Cre*^*M*^-mediated (as positive control) recombination in CD11b^-^CD45^-^ non-myeloid cells, microglia, and BAMs. N=3 mice per group.

Complementing the histological analysis, we employed flow cytometry on dissociated brain tissue to test the recombination activity of *Fcrls-2A-Cre* mice with an orthogonal approach. Specifically, we probed tdTomato expression in several populations, including non-myeloid cells, microglia, and BAMs that we gated based on reported surface markers (Figure 2N-P). While none of the non-myeloid cells were tdTomato-positive, we observed near complete tdTomato-positivity amongst microglia and BAMs in *Fcrls-2A-Cre; Ai14* and *Cx3cr1-Cre*^*M*^; *Ai14* mice which we used as a positive control (Figure 2Q-S). These findings collectively demonstrate that *Fcrls-2A-Cre* mice recombine floxed DNA sequences with high efficiency in all microglia and most BAMs.

### *Fcrls-2A-Cre* mice Efficiently Recombine Floxed DNA in Microglia Across Differentially Sensitive Cre-loxP Reporter Strains and in Early Development

The efficiency of Cre-mediated loxP site recombination and thus the completeness and fidelity depend on the expression levels of Cre and the distance between loxP sites amongst other factors (Glaser et al., 2005; Stifter and Greter, 2020). Previous carefully designed work using inducible microglia CreERT2 lines showed varying degrees of leakiness in the context of sensitive reporters and high CreERT2 expression levels (Chappell-Maor et al., 2020; Van Hove et al., 2020). Conversely, inefficient recombination has been observed in the context of less sensitive reporters and lower Cre expression levels (Alicia Bedolla et al., 2023; Travis E. Faust et al., 2023). To test the generalizability of loxP recombination by the Fcrls-2A-Cre line, we crossed *Fcrls-2A-Cre* mice to three different reporters of differential sensitivity and genomic context (Figure 3A-C). Amongst these reporters, *R26-Cas9-2A-GFP* and *R26-YFP* were created through insertion into the transcriptionally highly active *Rosa26* locus and driven by the strong *CAG* promoter, whereas *Rpl22-HA* is driven by its endogenous promoter and integrated in the endogenous *Rpl22* locus. Further, likely due to the long distance between the loxP sites, the *R26-YFP* reporter is known to be less sensitive compared to Ai14-type reporters and a good proxy for harder-to-recombine alleles. We co-immunostained brain slices and observed reporter activity in parenchymal IBA1-expressing microglia (Figure 3D-F). We found this activity in nearly all microglia (99%) and nearly all reporter signal (97-99%) was attributable to IBA1-expressing cells (Figure 3G-H). To further evaluate the completeness of recombination with an orthogonal flow cytometry assay, we dissociated and analyzed brains of *Fcrls-2A-Cre;R26-YFP* animals and noted that nearly all CD11b^+^CD45^lo^ microglia (98.7%) expressed YFP (Figure 3I-K). Together, these data indicate a generalizable and high level of completeness of recombination in *Fcrls-2A-Cre* mice independent of the reporter allele used.

**Figure 3.**
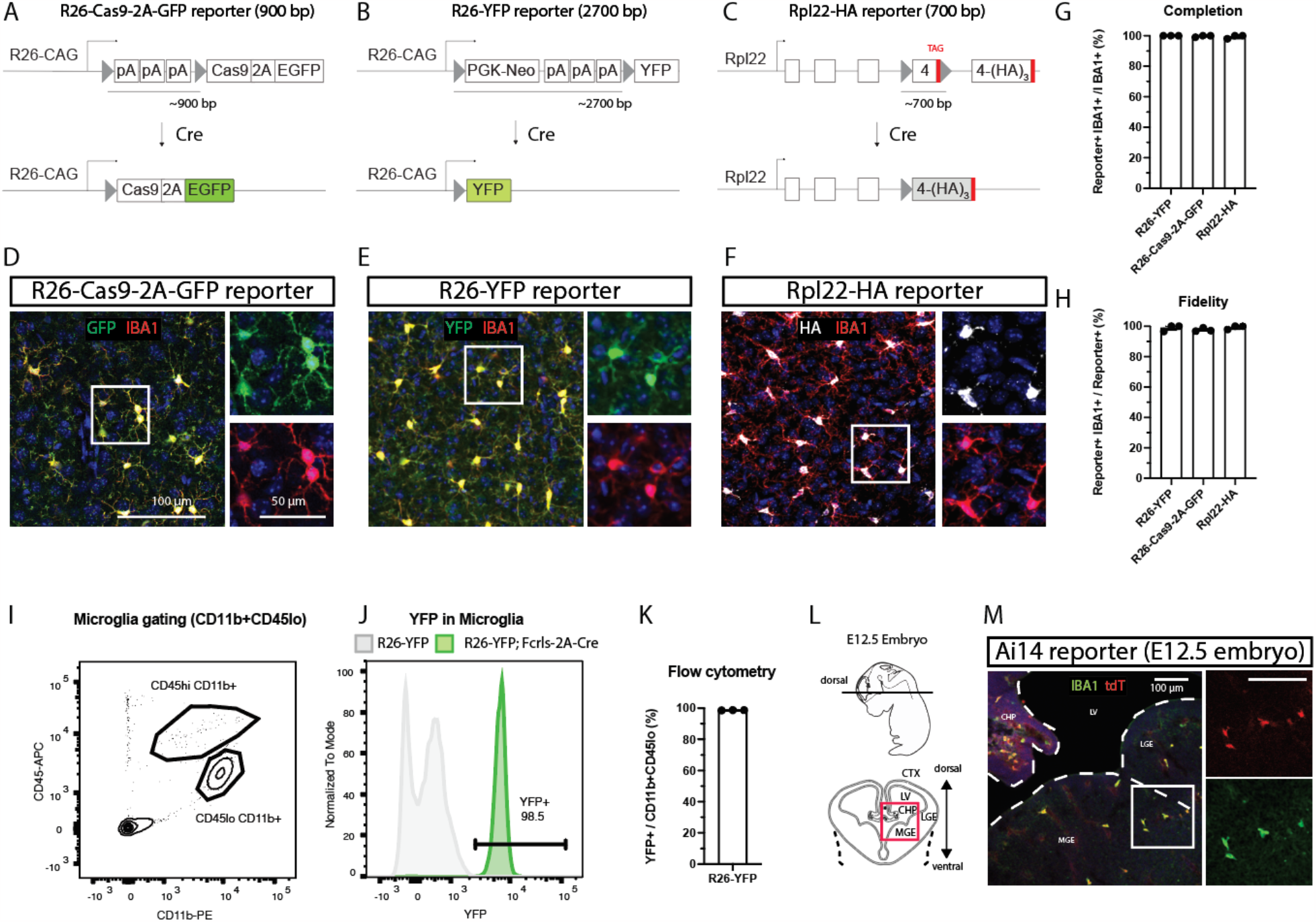
*Fcrls-2A-Cre* mice Efficiently Recombine Floxed DNA in Microglia Across Differentially Sensitive Cre-loxP Reporter Strains and in Early Development. **A-C**, Schematic representation of different Cre-loxP reporter stains employed. The different strains utilize different lox-stop reporter strategies and differ in the spacing between loxP sites, as well as the genomic context. Cre-mediated recombination of loxP sites results in expression of fluorescent reporters (EGFP, YFP) or an immunostainable HA-tagged ribosomal subunit (Rpl22-3xHA). **D-F**, Representative confocal micrographs of immunostained slices from *Fcrls-2A-Cre*^*+/-*^*;Reporter*^*+/-*^ mice showing GFP (green), YFP (green), or HA (white) detection possible due to Cre recombination in IBA1-immunostained microglia (red). **G-H**, Quantification of the completion and fidelity of Cre activity in IBA1^+^ parenchymal macrophages in the brain of the different reporter strains. N=3 mice per group. **I-J**, Representative flow cytometry density plot for microglia gating and histogram showing YFP expression in microglia. **K**, Quantification of YFP expression in microglia flow cytometry. N=3 mice. **L**, Schematic representation of E12.5 embryo, cryo-sectioning plane and imaging field of view (red box). CTX cortex, CHP choroid plexus, LGE lateral ganglionic eminence, LV lateral ventricle, MGE medium ganglionic eminence. **M**, Representative confocal micrograph of immunostained slices from *Fcrls-2A-Cre*^*+/-*^*;Ai14*^*+/-*^ mice showing Cre-dependent expression of tdTomato (red) and IBA1-positive microglia (green). Representative for N=2 E12.5 embryos.

Microglia-specific genes are not expressed uniformly and some genes express at lower levels or in select microglia subpopulations, especially during development (Hammond et al., 2019; Li et al., 2019), which could affect the utility of the *Fcrls-2A-Cre* line for studies requiring loxP recombination. To test recombination capability at embryonic development, we immunostained brain slices prepared from E12.5 *Fcrls-2A-Cre;Ai14* embryos (Figure 3L) and observed reporter activity in IBA1-expressing microglia, indicating that *Fcrls-2A-Cre* recombines floxed alleles at this early stage and is thus suitable for studies requiring early loxP recombination.

### *Fcrls-2A-Cre* mice Recombine Floxed DNA in a Subset of Peripheral Macrophages while Completely Sparing White Blood Cells

*Fcrls* is highly enriched in microglia compared to other myeloid cells (Butovsky et al., 2014), but even minute amounts of Cre can lead to all-or-none recombination events of floxed genomic sequences. To probe whether peripheral macrophages display Cre activity in *Fcrls-2A-Cre* mice, we examined liver and spleen tissue slices from *Fcrls-2A-Cre;R26-YFP* mice along with *Cx3cr1-Cre*^*M*^*;R26-YFP* mice as a positive control and comparator (Figure 4A). Across both organs, we observed near complete reporter activity (99% liver and 90% spleen) in tissues collected from *Cx3cr1-Cre*^*M*^*;R26-YFP* mice (Figure 4B-G). In contrast, samples from *Fcrls-2A-Cre;R26-YFP* mice showed substantially lower, albeit non-zero, recombination in these organs (23% liver and 14% spleen, Figure 4B-G), indicating a more modest targeting of peripheral macrophages, or perhaps only subsets thereof, in the *Fcrls-2A-Cre* line compared to the *Cx3cr1-Cre*^*M*^ line.

**Figure 4.**
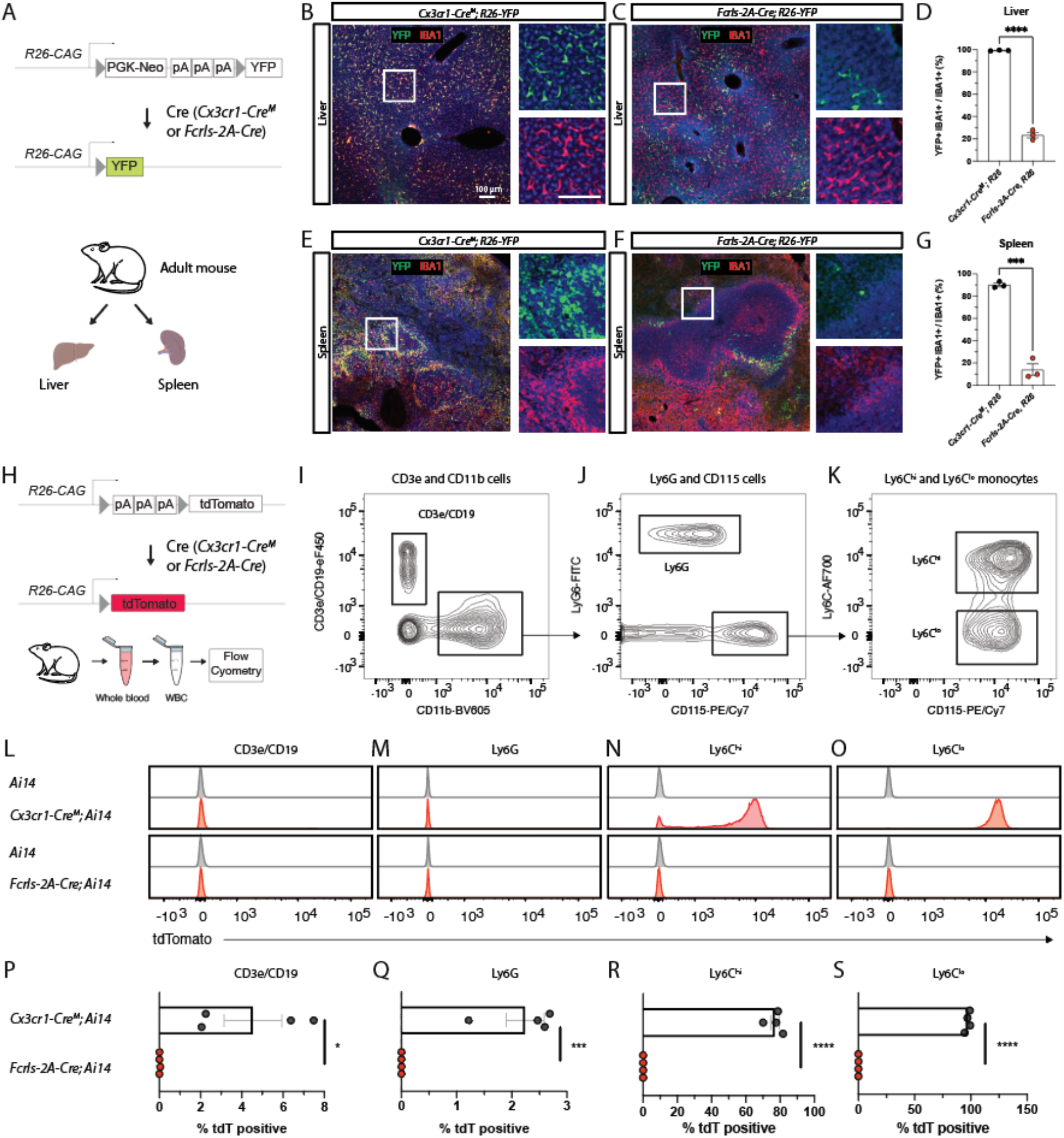
*Fcrls-2A-Cre* Mice Recombine Floxed DNA in A Subset of Peripheral Macrophages While Completely Sparing White Blood Cells. **A**, Schematic representation of recombination of R26-YFP reporter in crossings with *Fcrls-2A-Cre* and *Cx3cr1-Cre*^*M*^ (positive control) mice and organs harvested for immunostaining. **B-C**, Representative confocal micrographs of YFP fluorescence (green) and IBA1-immunostaining (red, macrophages) in the liver of adult animals. **D**, Quantification of YFP-positive macrophages in the liver. N=3 per group. Unpaired t-test. ***p<0.0001 **E-F**, Representative confocal micrographs of YFP fluorescence (green) and IBA1-immunostaining (red, macrophages) in the spleen of adult animals. **G**, Quantification of YFP-positive macrophages in the spleen. N=3 per group. Unpaired t-test. ***p<0.0002. **H**, Schematic representation of recombination of Ai14 reporter in crossings with *Fcrls-2A-Cre* and *Cx3cr1-Cre* (positive control) mice and white blood cell harvesting for flow cytometry. **I-K**, Representative flow cytometry density plots showing gating for lymphocytes (CD11b-CD19+CD3e+), granulocytes (CD11b+Ly6G+), and monocytes (CD115^+^Ly6C^hi^ and CD115^+^Ly6C^lo^). Pre-gated on single, live, CD45^+^ cells. **L-O**, Histograms showing tdTomato fluorescence among the different white blood cell subset populations for *Ai14* controls, *Cx3cr1-Cre*^*M*^; *Ai14* mice, and *Fcrls-Cre; Ai14* mice. **P-S**, Scatter plots showing the percentage of tdTomato-positive cells amongst all cells within a given subset in *Cx3cr1*^*M*^*-Cre*^*+/-*^*;Ai14*^*+/-*^ mice, and *Fcrls-2A-Cre*^*+/-*^*;Ai14*^*+/-*^ mice. N=4 mice per group. Unpaired t-test. *p=0.0178, ***p=0.0006, ****p<0.0001.

Monocytes infiltrate the brain and mediate pathological events in conditions such as stroke, Alzheimer’s disease, and multiple sclerosis (Henderson et al., 2009; Ritzel et al., 2015; Shang et al., 2016). Recently developed CreERT2 lines including *Tmem119-CreERT2* and *P2Ry12-CreERT2* almost completely avoid recombination of floxed alleles in monocytes (Kaiser and Feng, 2019; McKinsey et al., 2020). For constitutively active Cre lines, however, the currently best publicly available line, *Cx3cr1-Cre*^*M*^ (MMRRC, Gong et al., 2010; Zhao et al., 2019), is expected to express Cre in monocytes and perhaps other white blood cells based on the endogenous expression profile of *Cx3cr1* (ImmGen.org). This inability to discern the two cell types impedes unequivocal attribution of microglia function in these disorders when using this line. To determine whether *Fcrls-2A-Cre* displays Cre activity in leukocyte subsets, we harvested white blood cells from adult *Ai14* control mice, *Fcrls-2A-Cre; Ai14* mice or *Cx3cr1-2A-Cre*^*M*^; *Ai14* mice and stained for known markers (Figure 4H). Using flow cytometry, we gated single, live, CD45-expressing cells and further parsed them into CD3e/CD19^+^ lymphocytes, CD11b^+^Ly6G^+^ granulocytes, and CD115^+^Ly6C^hi^ and CD115^+^Ly6C^lo^ monocytes (Figure 4I-K). Histograms of tdTomato expression revealed that while different leukocyte subsets in *Cx3cr1-Cre*^*M*^; *Ai14* mice express tdTomato, virtually no leukocyte subsets from *Fcrls-2A-Cre; Ai14* mice do (Figure 4L-O). Specifically, quantification of tdTomato-positivity showed that none of the CD3e/CD19^+^ lymphocytes, CD11b^+^Ly6G^+^ granulocytes, CD115^+^Ly6C^hi^ monocytes, or CD115^+^Ly6C^lo^ monocytes expressed tdTomato in *Fcrls-2A-Cre;Ai14* mice compared to 4.5%, 2.2%, 77%, and 97% of these cells in *Cx3cr1-Cre*^*M*^; *Ai14* mice, respectively (Figure 4P-S). Together, these data demonstrate that, unlike the *Cx3cr1-Cre*^*M*^ line, the *Fcrls-2A-Cre* line avoids undesirable Cre recombinase activity in white blood cells, especially monocytes.

## Discussion

Transgenic mouse lines enabling the manipulation of floxed target genes in microglia are critical to advance our understanding of their biology. Additional publicly available, constitutively active Cre lines that spare white blood cells would be a great addition to our repertoire of mouse lines. In this study, we addressed this need by creating an *Fcrls-2A-Cre* mouse line. Using four different reporter strains, we demonstrate that the *Fcrls-2A-Cre* line effectively and specifically recombines floxed alleles in all microglia and most border associated macrophages (BAMs). Further we show that recombination occurs early in development and that a minority of tissue-resident macrophages in some peripheral tissues exhibit Cre activity as well. Finally, utilizing flow cytometric analysis, we demonstrate that Cre is virtually inactive in different white-blood cell subsets. Together, our studies show that *Fcrls-2A-Cre* is a powerful and easy-to-employ mouse line to study microglia while distinguishing them from white-blood cells. This new mouse line is immediately publicly available from JAX (Stock #036591).

*Fcrls* is one of the most highly expressed genes in microglia (Hammond et al., 2019), but its function remains unknown, since mice lacking *Fcrls* appear to be normal (Oleg Butovsky, unpublished data). To minimize potential knock-out-related issues, we chose a bicistronic knock-in approach using 2A peptide preserving endogenous expression and function of *Fcrls* (Figure 1A). Besides insertion of the exogenous *2A-Cre* sequence, development of the mice further required insertion of a point mutation at the PAM site of one of the CRISPR guide RNAs to prevent cutting of the targeting vector or targeted allele (Figure 1B). While the functional consequence of changing a single nucleotide in the 3’UTR is unknown and difficult to predict, the lack of conservation in rats, rabbits, and humans suggests that the change may be comparatively inconsequential. Supporting this notion, qPCR analysis showed that *Fcrls* mRNA expression was unaffected (Figure 1F). Moreover, *Fcrls-2A-Cre* mice were healthy, did not display any gross abnormalities, and have been bred to homozygosity (data not shown). Additionally, Cre expression in these mice did not perturb microglia phenotypes during early development (Figure 1F-J), a concern which has recently been reported for another microglia CreERT2 line (Sahasrabuddhe and Ghosh, 2022).

Our studies using *Ai14* reporter mice display that *Fcrls-2A-Cre* mice recombine floxed alleles efficiently and with high fidelity (Figure 2). The high completeness observed in microglia aligns well with *Fcrls* expression based on scRNA sequencing and *in-situ* hybridization studies (Hammond et al., 2019; Li et al., 2019). Also, in line with these sequencing studies, we observed recombination in a substantial fraction of border-associated macrophages (Figure 2F-I, S). Some, albeit generally much sparser, activity in some of these BAM subsets has been reported and can be inferred for the currently most specific inducible Cre lines *Tmem119-CreERT2, P2ry12-CreERT2*, and *Hexb-CreERT2* based on experimental data and endogenous expression pattern (Kaiser and Feng, 2019; Masuda et al., 2020; McKinsey et al., 2020). The continued difficulty to entirely avoid recombination in all BAM populations when using a highly potent Cre driver highlights a potential trade-off between completeness of recombination in microglia and distinction from BAMs. Researchers might find the inducible CreERT2 lines most suitable when specificity for microglia is the chief concern, whereas they might opt for constitutive Cre lines when completeness of recombination is the priority, particularly in the context of harder-to-recombine alleles. Yet another approach would be a successive use of Cre and CreERT2 in initial discovery and subsequent validation studies.

In this study, we evaluated Cre activity across four different Cre reporters and showed highly complete and specific combination in microglia, regardless of the reporter allele used (Figure 3A-K). This is of great significance given concerns for the efficiency of some CreERT2 systems, with several reports showing lower recombination efficiencies of harder-to-recombine alleles ranging from 40-70%, rarely reaching more than 90% even for the most potent CreERT2 lines that can come with a leakiness liability (Stifter and Greter, 2020; Alicia Bedolla et al., 2023; Travis E. Faust et al., 2023). The *Fcrls-2A-Cre* line recombined the reporter allele in 98% of microglia in the less sensitive R26-YFP reporter (Figure 3K), which indicates extremely robust activity. Ordinarily, lower activities may be perfectly acceptable for the study of reporter-labeled cells or genetic rescue by employing a STOP-flox rescue allele. However, in a common scenario with the goal of assessing a potential loss-of-function phenotype through knock-out of two, potentially harder-to-recombine endogenous alleles rather than through activation of an exogenous reporter, maximizing recombination efficiency is essential. This is a particular concern when testing for complex phenotypic changes that might require a near complete population-level knockout to precipitate the phenotype. *Fcrls-2A-Cre* mice should be a highly suitable tool for such applications.

With respect to temporal activity of Cre in *Fcrls-2A-Cre* mice, our studies at embryonic day 12.5 revealed recombinase activity at or even before this stage (Figure 3L-M), which renders this line suitable for studies during early development. To our knowledge, similar activity has not yet been shown for the *Cx3cr1-Cre*^*M*^ line and should not be assumed in absence of confirmation studies given available scRNA sequencing data that shows less consistent expression of this gene across several microglia subsets during development (Hammond et al., 2019; Li et al., 2019). Another line that does appear to recombine in these cells has recently been reported, named *Crybb1-Cre* (Brioschi et al., 2023). However, this line was not publicly available at the time of this study and has hence not been directly examined here.

Our examination of Cre activity in peripheral organs revealed that floxed alleles are recombined in a fraction of tissue-resident macrophages in the liver and spleen (Figure 4A-G). We observed recombination in a minority, yet non-negligible fraction, of these macrophages which was much lower than that seen in *Cx3cr1-Cre*^*M*^ mice. While recombination in even a subset of tissue-resident macrophages may appear surprising given the absence of *Fcrls* expression in adult peripheral macrophages (ImmGen.org), it is plausible based on transient *Fcrls* expression seen in some tissues at low to moderate levels during development (Mass et al., 2016). In such tissues with macrophages that turn over slowly, even low expression of Cre is sufficient for all-or-none recombination events. At this stage, we are unable to conclude if the labeled cells comprise a fraction of a general population or a nearly completely labeled specific subset of *Fcrls*-expressing macrophages. All in all, these data suggest a favorable recombination profile of *Fcrls-2A-Cre* over *Cx3cr1-Cre*^*M*^ in peripheral tissue-resident macrophage subsets, making it a suitable line when recombination in a minority of cells is acceptable and unlikely to produce a significantly confounding phenotype.

Finally, and most significantly, our flow cytometric studies show that Cre is virtually inactive in white blood cells (Figure 4H-S). This is of tremendous importance in the context of disease models since infiltration of monocytes is common in CNS diseases (Henderson et al., 2009; Ritzel et al., 2015; Shang et al., 2016). A frequently used work-around using inducible lines that display Cre activity in monocytes is to induce recombination with tamoxifen and wait for monocyte turnover, which is generally suitable for lineage tracing. However, the approach leaves ambiguity in disease contexts where monocytes with recombined alleles may at least partially drive pathology before getting turned over. Compared to other lines, *Fcrls-2A-Cre* mice offer unique advantages. The currently best publicly available constitutively active Cre line *Cx3cr1-Cre*^*M*^ labels a large fraction of monocytes and a small fraction of lymphocytes and granulocytes (Figure 4P-S) and other lines that achieve similar specificity, namely *P2ry12-CreERT2* and *Tmem119-CreERT2*, require tamoxifen administration and are likely less efficient at recombining the allele in microglia (Kaiser and Feng, 2019; McKinsey et al., 2020). Taken together, our observation that *Fcrls-2A-Cre* mice spare monocytes and other white blood cells makes this novel line extraordinarily suitable for studies where constitutive activity of Cre is desired and where monocytes would be a major confounding factor.

*Fcrls-2A-Cre* mice present an important addition to the microglia toolbox. In the future, more and even better constitutively active Cre lines may be developed. Most recently, *Cx3cr1* and *Sall1* loci were harnessed to create a split Cre mouse line that displayed impressive fidelity (Kim et al., 2021). At the same time, an obstacle to implementing this line for knockout studies is its multiallelic design and relative inefficiency that may require breeding of two split-Cre loci to homozygosity in addition to the homozygous floxed allele. Perhaps, another combination for a split Cre approach achieving similar specificity while retaining high efficiency could involve targeting *P2ry12* and *Fcrls* on the same allele on chromosome 3. This would facilitate breeding and genetic crosses of the mice. Moving forward, combining these improved mouse lines with emerging strategies to deliver genetic material to microglia through AAV and other means (Lin et al., 2022; Young et al., 2023) will facilitate major advances in research into microglia function in health and disease.

## References

Aguzzi A, Zhu C (2017) Microglia in prion diseases. The Journal of clinical investigation 127:3230–3239.

Alicia Bedolla, Gabriel Mckinsey, Kierra Ware, Nicolas Santander, Thomas Arnold, Yu Luo (2023) Finding the right tool: a comprehensive evaluation of microglial inducible cre mouse models. bioRxiv:2023.04.17.536878.

Bennett ML, Bennett FC (2020) The influence of environment and origin on brain resident macrophages and implications for therapy. Nature Neuroscience 23:157–166.

Bennett ML, Bennett FC, Liddelow SA, Ajami B, Zamanian JL, Fernhoff NB, Mulinyawe SB, Bohlen CJ, Adil A, Tucker A (2016) New tools for studying microglia in the mouse and human CNS. Proceedings of the National Academy of Sciences 113:E1738–E1746.

Brioschi S, Belk JA, Peng V, Molgora M, Rodrigues PF, Nguyen KM, Wang S, Du S, Wang W-L, Grajales-Reyes GE, Ponce JM, Yuede CM, Li Q, Baer JM, DeNardo DG, Gilfillan S, Cella M, Satpathy AT, Colonna M (2023) A Cre-deleter specific for embryo-derived brain macrophages reveals distinct features of microglia and border macrophages. Immunity 56:1027–1045.e8.

Butovsky O, Jedrychowski MP, Moore CS, Cialic R, Lanser AJ, Gabriely G, Koeglsperger T, Dake B, Wu PM, Doykan CE (2014) Identification of a unique TGF-β–dependent molecular and functional signature in microglia. Nature neuroscience 17:131–143.

Chappell-Maor L, Kolesnikov M, Kim J-S, Shemer A, Haimon Z, Grozovski J, Boura-Halfon S, Masuda T, Prinz M, Jung S (2020) Comparative analysis of CreER transgenic mice for the study of brain macrophages: A case study. European Journal of Immunology 50:353–362.

Clausen B, Burkhardt C, Reith W, Renkawitz R, Förster I (1999) Conditional gene targeting in macrophages and granulocytes using LysMcre mice. Transgenic research 8:265–277.

Davalos D, Grutzendler J, Yang G, Kim JV, Zuo Y, Jung S, Littman DR, Dustin ML, Gan W-B (2005) ATP mediates rapid microglial response to local brain injury in vivo. Nature neuroscience 8:752–758.

Dumas AA, Borst K, Prinz M (2021) Current tools to interrogate microglial biology. Neuron 109:2805–2819.

Ferron M, Vacher J (2005) Targeted expression of Cre recombinase in macrophages and osteoclasts in transgenic mice. Genesis 41:138–145.

Glaser S, Anastassiadis K, Stewart AF (2005) Current issues in mouse genome engineering. Nature Genetics 37:1187–1193.

Gong S, Kus L, Heintz N (2010) Rapid bacterial artificial chromosome modification for large-scale mouse transgenesis. Nature protocols 5:1678–1696.

Hagemeyer N, Hanft K-M, Akriditou M-A, Unger N, Park ES, Stanley ER, Staszewski O, Dimou L, Prinz M (2017) Microglia contribute to normal myelinogenesis and to oligodendrocyte progenitor maintenance during adulthood. Acta neuropathologica 134:441–458.

Haimon Z, Volaski A, Orthgiess J, Boura-Halfon S, Varol D, Shemer A, Yona S, Zuckerman B, David E, Chappell-Maor L (2018) Re-evaluating microglia expression profiles using RiboTag and cell isolation strategies. Nature immunology 19:636–644.

Hammond TR, Dufort C, Dissing-Olesen L, Giera S, Young A, Wysoker A, Walker AJ, Gergits F, Segel M, Nemesh J (2019) Single-cell RNA sequencing of microglia throughout the mouse lifespan and in the injured brain reveals complex cell-state changes. Immunity 50:253–271.

Henderson AP, Barnett MH, Parratt JD, Prineas JW (2009) Multiple sclerosis: distribution of inflammatory cells in newly forming lesions. Annals of Neurology: Official Journal of the American Neurological Association and the Child Neurology Society 66:739–753.

Itagaki S, McGeer P, Akiyama H, Zhu S, Selkoe D (1989) Relationship of microglia and astrocytes to amyloid deposits of Alzheimer disease. Journal of neuroimmunology 24:173–182.

Kaiser T, Feng G (2019) Tmem119-EGFP and Tmem119-CreERT2 transgenic mice for labeling and manipulating microglia. eneuro 6.

Keren-Shaul H, Spinrad A, Weiner A, Matcovitch-Natan O, Dvir-Szternfeld R, Ulland TK, David E, Baruch K, Lara-Astaiso D, Toth B (2017) A unique microglia type associated with restricting development of Alzheimer’s disease. Cell 169:1276–1290.

Kim J-S, Kolesnikov M, Peled-Hajaj S, Scheyltjens I, Xia Y, Trzebanski S, Haimon Z, Shemer A, Lubart A, Van Hove H (2021) A binary Cre transgenic approach dissects microglia and CNS border-associated macrophages. Immunity 54:176–190.

Kim W-K, Alvarez X, Fisher J, Bronfin B, Westmoreland S, McLaurin J, Williams K (2006) CD163 identifies perivascular macrophages in normal and viral encephalitic brains and potential precursors to perivascular macrophages in blood. The American journal of pathology 168:822–834.

Kisanuki YY, Hammer RE, Miyazaki J, Williams SC, Richardson JA, Yanagisawa M (2001) Tie2-Cre transgenic mice: a new model for endothelial cell-lineage analysis in vivo. Developmental biology 230:230–242.

Lawson LJ, Perry VH, Dri P, Gordon S (1990) Heterogeneity in the distribution and morphology of microglia in the normal adult mouse brain. Neuroscience 39:151–170.

Li Q, Cheng Z, Zhou L, Darmanis S, Neff NF, Okamoto J, Gulati G, Bennett ML, Sun LO, Clarke LE (2019) Developmental heterogeneity of microglia and brain myeloid cells revealed by deep single-cell RNA sequencing. Neuron 101:207–223.

Lin R, Zhou Y, Yan T, Wang R, Li H, Wu Z, Zhang X, Zhou X, Zhao F, Zhang L, Li Y, Luo M (2022) Directed evolution of adeno-associated virus for efficient gene delivery to microglia. Nature Methods 19:976–985.

Madisen L, Zwingman TA, Sunkin SM, Oh SW, Zariwala HA, Gu H, Ng LL, Palmiter RD, Hawrylycz MJ, Jones AR (2010) A robust and high-throughput Cre reporting and characterization system for the whole mouse brain. Nature neuroscience 13:133–140.

Mass E, Ballesteros I, Farlik M, Halbritter F, Günther P, Crozet L, Jacome-Galarza CE, Händler K, Klughammer J, Kobayashi Y (2016) Specification of tissue-resident macrophages during organogenesis. Science 353.

Masuda T, Amann L, Sankowski R, Staszewski O, Lenz M, Snaidero N, Jordão MJC, Böttcher C, Kierdorf K, Jung S (2020) Novel Hexb-based tools for studying microglia in the CNS. Nature immunology 21:802–815.

Mathys H, Adaikkan C, Gao F, Young JZ, Manet E, Hemberg M, De Jager PL, Ransohoff RM, Regev A, Tsai L-H (2017) Temporal tracking of microglia activation in neurodegeneration at single-cell resolution. Cell reports 21:366–380.

McKinsey GL, Lizama CO, Keown-Lang AE, Niu A, Santander N, Larpthaveesarp A, Chee E, Gonzalez FF, Arnold TD (2020) A new genetic strategy for targeting microglia in development and disease. Elife 9:e54590.

Miron VE, Priller J (2020) Investigating Microglia in Health and Disease: Challenges and Opportunities. Trends in Immunology 41:785–793.

O’Loughlin E, Madore C, Lassmann H, Butovsky O (2018) Microglial phenotypes and functions in multiple sclerosis. Cold Spring Harbor perspectives in medicine 8:a028993.

Orthgiess J, Gericke M, Immig K, Schulz A, Hirrlinger J, Bechmann I, Eilers J (2016) Neurons exhibit Lyz2 promoter activity in vivo: Implications for using LysM-Cre mice in myeloid cell research. European journal of immunology 46:1529–1532.

Paolicelli RC, Bolasco G, Pagani F, Maggi L, Scianni M, Panzanelli P, Giustetto M, Ferreira TA, Guiducci E, Dumas L (2011) Synaptic pruning by microglia is necessary for normal brain development. science 333:1456–1458.

Parkhurst CN, Yang G, Ninan I, Savas JN, Yates III JR, Lafaille JJ, Hempstead BL, Littman DR, Gan W-B (2013) Microglia promote learning-dependent synapse formation through brain-derived neurotrophic factor. Cell 155:1596–1609.

Ransohoff RM (2016) How neuroinflammation contributes to neurodegeneration. Science 353:777–783.

Ritzel RM, Patel AR, Grenier JM, Crapser J, Verma R, Jellison ER, McCullough LD (2015) Functional differences between microglia and monocytes after ischemic stroke. Journal of neuroinflammation 12:1–12.

Sahasrabuddhe V, Ghosh HS (2022) Cx3Cr1-Cre induction leads to microglial activation and IFN-1 signaling caused by DNA damage in early postnatal brain. Cell reports 38:110252.

Samokhvalov IM, Samokhvalova NI, Nishikawa S (2007) Cell tracing shows the contribution of the yolk sac to adult haematopoiesis. Nature 446:1056–1061.

Schafer DP, Lehrman EK, Kautzman AG, Koyama R, Mardinly AR, Yamasaki R, Ransohoff RM, Greenberg ME, Barres BA, Stevens B (2012) Microglia sculpt postnatal neural circuits in an activity and complement-dependent manner. Neuron 74:691–705.

Shang DS, Yang YM, Zhang H, Tian L, Jiang JS, Dong YB, Zhang K, Li B, Zhao WD, Fang WG (2016) Intracerebral GM-CSF contributes to transendothelial monocyte migration in APP/PS1 Alzheimer’s disease mice. Journal of Cerebral Blood Flow & Metabolism 36:1978–1991.

Sierra A, Encinas JM, Deudero JJ, Chancey JH, Enikolopov G, Overstreet-Wadiche LS, Tsirka SE, Maletic-Savatic M (2010) Microglia shape adult hippocampal neurogenesis through apoptosis-coupled phagocytosis. Cell stem cell 7:483–495.

Stevens B, Allen NJ, Vazquez LE, Howell GR, Christopherson KS, Nouri N, Micheva KD, Mehalow AK, Huberman AD, Stafford B (2007) The classical complement cascade mediates CNS synapse elimination. Cell 131:1164–1178.

Stifter SA, Greter M (2020) STOP floxing around: Specificity and leakiness of inducible Cre/loxP systems. European Journal of Immunology 50:338–341.

Travis E. Faust, Philip A. Feinberg, Ciara O’Connor, Riki Kawaguchi, Andrew Chan, Haley Strasburger, Takahiro Masuda, Lukas Amann, Klaus-Peter Knobeloch, Marco Prinz, Anne Schaefer, Dorothy P. Schafer (2023) A comparative analysis of microglial inducible Cre lines. bioRxiv:2023.01.09.523268.

Ueno M, Fujita Y, Tanaka T, Nakamura Y, Kikuta J, Ishii M, Yamashita T (2013) Layer V cortical neurons require microglial support for survival during postnatal development. Nature neuroscience 16:543–551.

Van Hove H, Antunes ARP, De Vlaminck K, Scheyltjens I, Van Ginderachter JA, Movahedi K (2020) Identifying the variables that drive tamoxifen-independent CreERT2 recombination: Implications for microglial fate mapping and gene deletions. European Journal of Immunology 50:459–463.

Weinhard L, Di Bartolomei G, Bolasco G, Machado P, Schieber NL, Neniskyte U, Exiga M, Vadisiute A, Raggioli A, Schertel A (2018) Microglia remodel synapses by presynaptic trogocytosis and spine head filopodia induction. Nature communications 9:1–14.

Wieghofer P, Prinz M (2016) Genetic manipulation of microglia during brain development and disease. Biochimica et Biophysica Acta (BBA)-Molecular Basis of Disease 1862:299–309.

Wlodarczyk A, Holtman IR, Krueger M, Yogev N, Bruttger J, Khorooshi R, Benmamar-Badel A, de Boer-Bergsma JJ, Martin NA, Karram K (2017) A novel microglial subset plays a key role in myelinogenesis in developing brain. The EMBO journal 36:3292–3308.

Yona S, Kim K-W, Wolf Y, Mildner A, Varol D, Breker M, Strauss-Ayali D, Viukov S, Guilliams M, Misharin A (2013) Fate mapping reveals origins and dynamics of monocytes and tissue macrophages under homeostasis. Immunity 38:79–91.

Young A, Neumann B, Segel M, Chen CZ-Y, Tourlomousis P, Franklin RJ (2023) Targeted evolution of adeno-associated virus capsids for systemic transgene delivery to microglia and tissue-resident macrophages. Proceedings of the National Academy of Sciences 120:e2302997120.

Zhao X-F, Alam MM, Liao Y, Huang T, Mathur R, Zhu X, Huang Y (2019) Targeting microglia using Cx3cr1-Cre lines: revisiting the specificity. eneuro 6.

